# Deep Learning Enhanced Tandem Repeat Variation Identification via Multi-Modal Conversion of Nanopore Reads Alignment

**DOI:** 10.1101/2023.08.17.553659

**Authors:** Xingyu Liao, Juexiao Zhou, Bin Zhang, Xiaopeng Xu, Haoyang Li, Xin Gao

## Abstract

Identification of tandem repeat (TR) variations plays a crucial role in advancing our understanding of genetic diseases, forensic analysis, evolutionary studies, and crop improvement, thereby contributing to various fields of research and practical applications. However, traditional TR identification methods are often limited to processing genomes obtained through sequence assembly and cannot directly start detection from sequencing reads. Furthermore, the inflexibility of detection mode and parameters hinders the accuracy and completeness of the identification, rendering the results unsatisfactory. These shortcomings result in existing TR variation identification methods being associated with high computational cost, limited detection sensitivity, precision and comprehensiveness. Here, we propose DeepTRs, a novel method for identifying TR variations, which enables direct TR variation identification from raw Nanopore sequencing reads and achieves high sensitivity, accuracy, and completeness results through the multi-modal conversion of Nanopore reads alignment and deep learning. Comprehensive evaluations demonstrate that DeepTRs outperform existing methods.

## 1 Introduction

Tandem repeats (TRs) are repetitive DNA sequences that occur adjacent to each other in a head-to-tail manner within a genome [1]. They are characterized by consecutive repetitions of a specific short nucleotide sequence motif. TRs can vary in length and the number of repetitions, making them highly polymorphic across individuals and species [2]. Expansions of TRs refer to the phenomenon where the number of repetitions in a TR region increases in subsequent generations [3]. These expansions can occur due to replication slippage, a mechanism during DNA replication where errors lead to the addition or deletion of repeat units. As a result, the length of the tandem repeat region increases, leading to instability in the genome. Expansions of certain TR regions have been associated with various genetic disorders [4]. For example, the expansion of trinucleotide repeat sequences, such as CAG or CGG repeats, have been linked to disorders including cancer [5], ASD [6], Huntington’s disease [7], fragile X syndrome [8], and myotonic dystrophy [9]. These expansions can disrupt gene function and contribute to the development of specific diseases or disorders.

TR identification methods play a crucial role in deciphering the repetitive nature of DNA sequences within genomes. These methods employ diverse computational techniques to detect and analyze regions characterized by the repetition of specific DNA motifs. These methods can be subdivided into the following six types: 1) Periodicity-based methods, which utilize algorithms such as Fourier transform or sequence alignment to identify the periodic structure of tandem repeats. Tandem Repeats Finder (TRF) [10], RepeatMasker [11], XSTREAM [12], and T-REKS [13] are typical representatives of periodicity-based methods. 2) Seed-and-Extend approaches, which iteratively extend seed motifs to detect TR regions with variable lengths and repetitions. Sputnik [14], TROLL [15], TRUST [16], and TRstalker [17] are examples of seed-and-extend approaches. 3) Unique k-mer counting methods, which leverage the overrepresentation of specific k-mers to locate TR regions. Tandem Repeats Database (TRDB) [18], GMATo [19], and IDR [20] are commonly used tools for seed-and-extend strategies in TR identification. 4) Graph-based methods, which construct graphs to capture the repetitive structure and identify TRs through graph analysis. Tandem Repeat Graph Finder (TRGF) [21], TRAP [22], STR-FM [23], and Tandem Repeats Graph Comparison (TRGC) [24] are examples of graph-based methods used for TR identification. 5) Machine learning-based methods, which employ various algorithms to classify TR regions based on sequence features. The combined utilization of these approaches facilitates comprehensive identification and characterization of TRs, aiding in the understanding of their functional and evolutionary roles in genomes. PopAffiliator [25], Tally-2.0 [26], RExPRT [27], and WarpSTR [28] are TR identification methods developed based on machine learning. 6) Deep learning-based methods, which based on deep learning typically involve training models to recognize patterns found in TRs. DeepRepeat [29], DeepSymmetry [30], NanoSTR [31] are representative methods based on deep learning.

The TR identification methods summarized above are often limited to processing genomes obtained through sequence assembly and cannot directly start detection from sequencing reads. Furthermore, the inflexibility of detection mode and parameters hinders the accuracy and completeness of the identification, rendering the results unsatisfactory. In recent years, with the rapid development of deep learning technology, it has made significant contributions to bioinformatics, revolutionizing many areas of genomic analysis, including gene expression, protein structure prediction, variant calling, and functional genomics [32]. Multimodal conversion refers to the integration of diverse types of biological data or modalities, such as genomic, epigenomic, transcriptomic, proteomic, or imaging data [33], [34]. Integration of multiple data modalities provides a more comprehensive and holistic view of biological systems. It enables researchers to capture different layers of information and relationships between various biological components, leading to a deeper understanding of complex biological processes. However, there is no specific method based on multi-modal conversion combined with deep learning that has been proposed for TR identification so far.

TR expansion identification relies heavily on accurate TR identification as a foundational step. The process of detecting TR expansions involves identifying the baseline length and structure of the TRs in a given genome or sequence. Once accurate TR identification has been achieved, the subsequent step involves comparing the identified TRs to other samples or reference genomes to detect any variations in repeat length or copy number. By comparing the known TR structures to the sequences under examination, potential expansions or contractions can be inferred. There are four common classes of tools used for identifying tandem repeat expansions: 1) Southern Blotting, which is a classical molecular biology technique used to detect TR expansions. It involves DNA digestion, gel electrophoresis, DNA transfer to a membrane, and hybridization with a labeled probe targeting the TR region of interest [35]. Repeat expansion can be identified by the presence of extra bands or smears on the blot. 2) PCR-based methods, which utilizes primers designed to specifically target the TR region flanking sequences and employs a denaturation step that selectively amplifies expanded alleles. Triplet Repeat-Primed PCR (TPPCR) [36] is a modification of RP-PCR for specifically detecting trinucleotide repeat expansions. 3) NGS-based methods, which scans the genomic sequences for TR motifs, quantifies the repeat sizes, and identifies potential expansions from NGS data. ExpansionHunter [37], ExpansionHunter Denovo [38], and STRetch [39] are typical examples of NGS-based methods. 4) Single-Molecule sequencing-based methods, which enables direct detection of expanded alleles form long reads without PCR, providing accurate and comprehensive assessment of repeat size and structure. tandem-genotypes [40], NanoSatellite [41], TRiCoLOR [42], and Straglr [43] are methods for identifying TR expansions based on long reads leverage the capabilities of long-read sequencing technologies, such as Oxford Nanopore and Pacific Biosciences (PacBio) sequencing.

Indeed, many existing algorithms for TR expansion detection still heavily rely on traditional identification strategies introduced above, which can be associated with high computational costs, limited detection sensitivity, precision, and comprehensiveness. These challenges highlight the need for further advancements in TR expansion detection methods to overcome these limitations and improve the accuracy and efficiency of TR expansion identification. In this study, we proposed DeepTRs, a novel method for identifying TR expansions and even variations, which enables direct TR variation identification from raw Nanopore sequencing reads and achieves high sensitivity, accuracy, and completeness results through the multi-modal conversion of Nanopore reads alignment and deep learning. Comprehensive evaluations demonstrate that DeepTRs outperform existing methods.

## 2 Results

### 2.1 Generation of input data for DeepTRs

#### 2.1.1 Generating ground-truth

In this study, we firstly collected the assemblies of samples HG001, HG002, HG003, HG004, HG005, HG00733, and human reference genome (GRCh38.p13) from NCBI. Secondly, two most famous TRs identification tools TRF (https://tandem.bu.edu/trf/trf.html) and Repeatmasker (https://www.repeatmasker.org/) are used to determine the TR regions on the selected assemblies and reference genome. Finally, the detection results of TRF and Repeatmasker on each dataset are combined to form the ground-truth for each dataset.

#### 2.1.2 Mapping nanopore reads to assembly and the human reference genome

The multiple sequence alignment tool minimap2 [44] is employed to map the nanopore sequencing raw reads of samples HG001, HG002, HG003, HG004, HG005, and HG00733 to the respective assembly. Additionally, it also used to map the nanopore raw reads from lung cancer samples DRR171452, DRR171453, DRR171454, DRR171429, DRR171430, DRR171431, DRR171432, DRR171433, and DRR203145 to the human reference genome.

#### 2.1.3 Generating alignment regions around the ground truth

The samtools [45] is used to generate the initial alignment regions based on the ground-truth determined in section 2.1.1 and the output of minimap2 obtained in section 2.1.2. Next, the final alignment regions are obtained by extending the left and right boundaries of the initial alignment regions to both ends by 5 times the length of the initial alignment regions. After that, the final alignment regions will be used as the input data for DeepTRs. Actually, the initial alignment region refers to TR region identified through the joint efforts of TRF and RepeatMasker. Conversely, the final alignment region encompasses the region used for detecting potential variations, as we hypothesize that mutations are likely to occur within the expansion region.

### 2.2 Evaluation of the TR identification of DeepTRs

To evaluate the performance of DeepTRs in TR identification, we compared the detected motifs, boundaries, and range size with the ground-truth. In this evaluation, the nanpore sequencing reads of human samples HG001, HG002, HG003, HG004, HG005 and HG00733, and their corresponding assemblies are used to measure the accuracy of the TR identification of DeepTRs. Firstly, we evaluated the precision and recall of TR identification results of DeepTRs based on the ground truth regions. According to Fig.2(D), the overall precision reaches 0.88, while the recall reaches 0.98, indicating that 98% of ground-truth regions can be covered by the detection results of DeepTRs. Additionally, we conducted tests on the boundary difference between the detected regions of DeepTRs and the ground-truth regions. As shown in Fig.2(E), the distribution of the boundary difference has a good enrichment effect near 0, and over 80% of difference are within 50 bp. We also compared the performance of DeepTRs with some traditional TR detection tools, such as T-REKS, mreps and TRASH. The detailed experimental results are displayed in Section S2 of the supplementary. The evaluation results of those experiments show that DeepTRs can identify the TR regions marked in the ground-truth with high integrity and accuracy, which also proves the effectiveness of the adopted models.

**Fig. 1.**
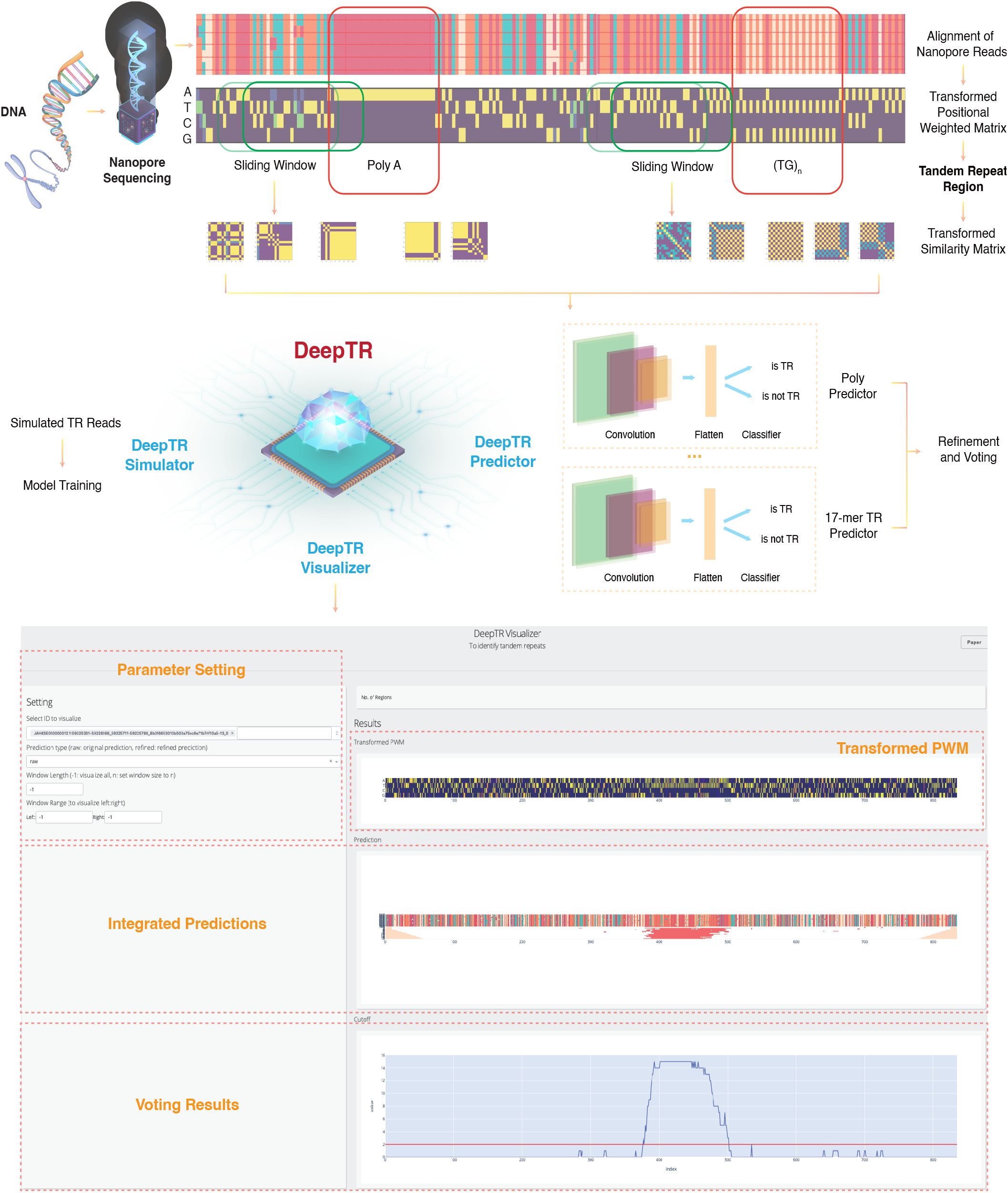
The illustration of DeepTR. DeepTR is composed of three main components: the DeepTR Simulator, the DeepTR Predictor, and the DeepTR Visualizer. Nanopore sequencing is employed to sequence the DNA, and the resulting nanopore reads are aligned and transformed into a positional weighted matrix (PWM). Subsequently, DeepTR converts the PWM into transformed similarity matrices (TSM) using modal conversion, which serve as inputs for the DeepTR Predictor. The DeepTR Predictor utilizes 17 deep learning models trained with the simulated data from the DeepTR Simulator in parallel to predict the probability of 1-17mer TRs individually, and the final prediction for the base corresponding to the middle position of the sliding window is determined through a voting process. The prediction results can be conveniently visualized using the DeepTR Visualizer.

**Fig. 2.**
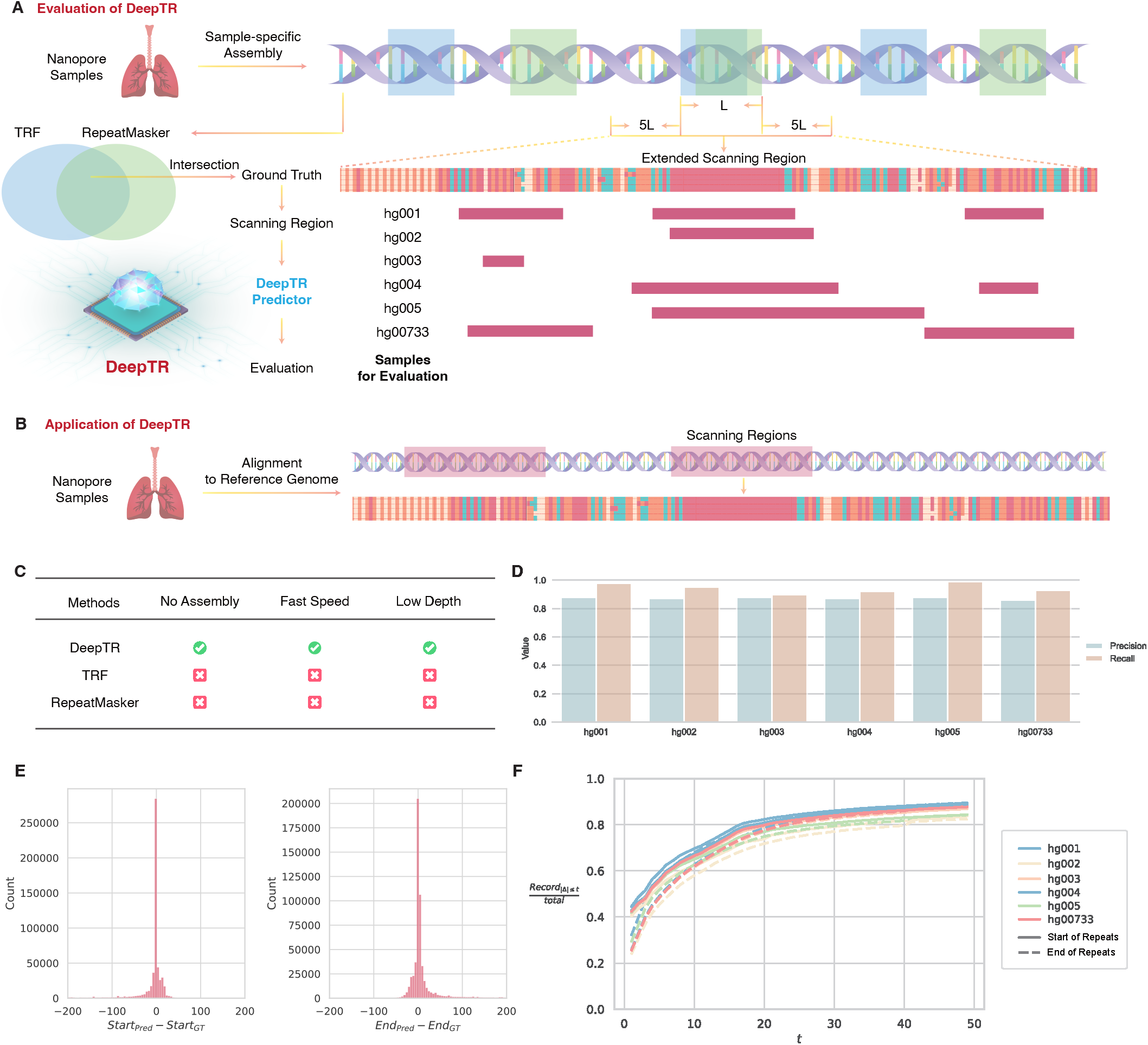
Evaluation of the TR identification of DeepTRs. A) the principle of generating ground-truth and extracting intervals for DeepTRs scanning. B) the application scenario of DeepTRs. C) the obvious advantages of DeepTRs compared with the two most famous TR identification tools TRF and RepeatMasker. D) the overall precision and recall of identification results generated by DeepTRs on six datasets. E) the boundary difference between the detected regions of DeepTRs and the ground-truth regions.

### 2.3 Evaluation of the TR variation identification of DeepTRs

The ultimate purpose of the proposed algorithm is to identify TR variations present in the genome of the sequenced sample. In this study, while assessing the accuracy of the model, the assembly results corresponding to the sequencing samples are used as the reference genomes. However, when conducting the TR expansions assay, the human genome hg38 is utilized as the reference. As described before, the tools TRF and Repeatmasker are used to determine the TR regions on the human reference genome (hg38), and the their detection results are combined to form the groundtruth. After that, the nanpore sequencing reads are mapped to the human reference genome by using minimap2. When scanning the potential TR interval, DeepTR will expand to the left and right ends by 5 times its own length according to the boundary of the interval in the ground truth. Furthermore, when the TR interval is determined, their left and right boundaries will be compared with the boundaries of the corresponding region in the ground-truth. If the boundary difference is larger than 20 bp, the detected TR interval will be considered as the potential TR variation interval. The principle of identifying TR variations is shown in Fig.3(A).

To evaluate the TR variation identification performance of DeepTRs, we compared the effect of DeepTR with two recently proposed tools Straglr and RExPRT. The evaluation results presented in Section S3 of the supplementary materials demonstrate that DeepTRs can accurately and sensitively identify TR variations, highlighting the superiority of the adopted models. Specifically, in Figure 3(B), the groundtruth region spans from position 26,517,115 to 26,517,144 on the chromosome NC 000021.9. However, DeepTRs identify the TR region boundaries as 26,517,114 and 26,517,171, respectively. Both the consensus sequences of the groundtruth region and the region identified by DeepTRs consist of a high-density sequence of the character ‘A’. Notably, during alignment to the human reference genome, soft clipping is observed in the CIGAR string for the consensus sequence obtained by DeepTRs. This suggests the presence of sequence fragments in the consensus sequence that are not present in the reference genome, indicating the existence of TR variations.

**Fig. 3.**
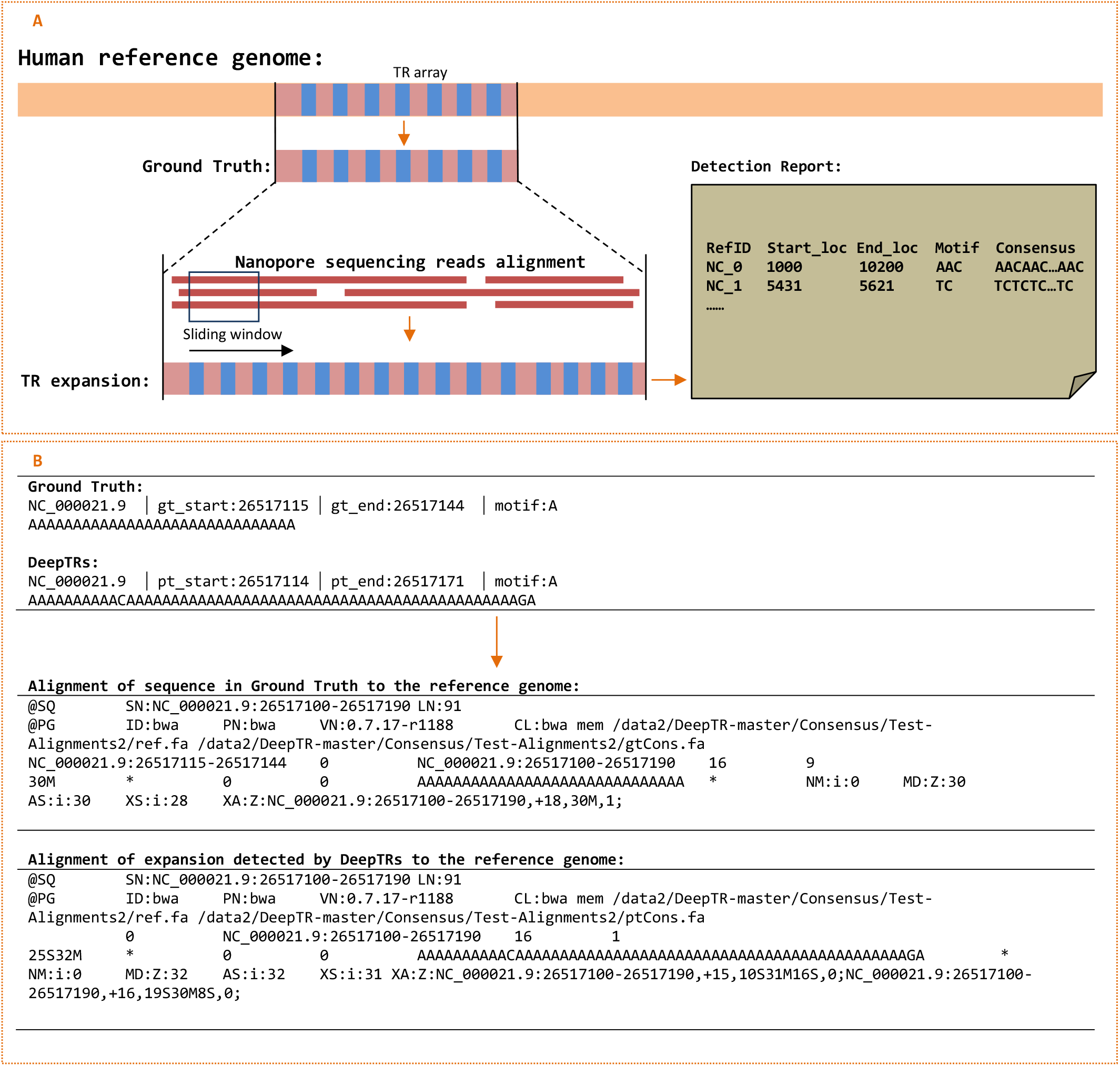
The principle and evaluation of DeepTRs’s TR variation identification. A) the principle of TR variation identification in DeepTRs. B) A concrete example of TR extensions discovered by DeepTRs. The aligner and the reference genome used in B) are bwa and hg38, repectively. The CIGAR string is how the SAM/BAM format represents spliced alignments.

## 3 Methods

### 3.1 TR simulation for model training with DeepTR Simulator

We developed a tandem repeat simulator (TRS) to simulate the nanopore reads containing tandem repeats (Algorithm 1). Since tandem repeats do not have sequence motifs, we only need to simulate sequences with repetitive structure features for model training. TRS allows users to specify the length of the generated sequence and the length of the repeat region. The size of the repeat unit (*k*) could be selected from 1 to 17, and the number of repeat units can also be set. Based on TRS, we could generate a batch of sequences with a TR in the middle and non-TR regions at both ends. Then, TRS simulates mutations, insertion, and deletion to introduce errors into those sequences to simulate nanopore reads containing tandem repeats.

For each *k*-mer tandem repeat, we kept the values of *p*_*i*_, *p*_*d*_, and *p*_*m*_ fixed at 0.05 and randomly chose a value of *t* between 5 to 100. We generated 10,000 sequences with these parameters. Then, we used a sliding window with a length of 7*k* to generate sub-sequences for each generated region. We labeled sub-sequences with intermediate positions in the tandem repeat region as Positive and sub-sequences with intermediate positions in the non-tandem repeat region as Negative.

#### Algorithm 1 DeepTR Simulator

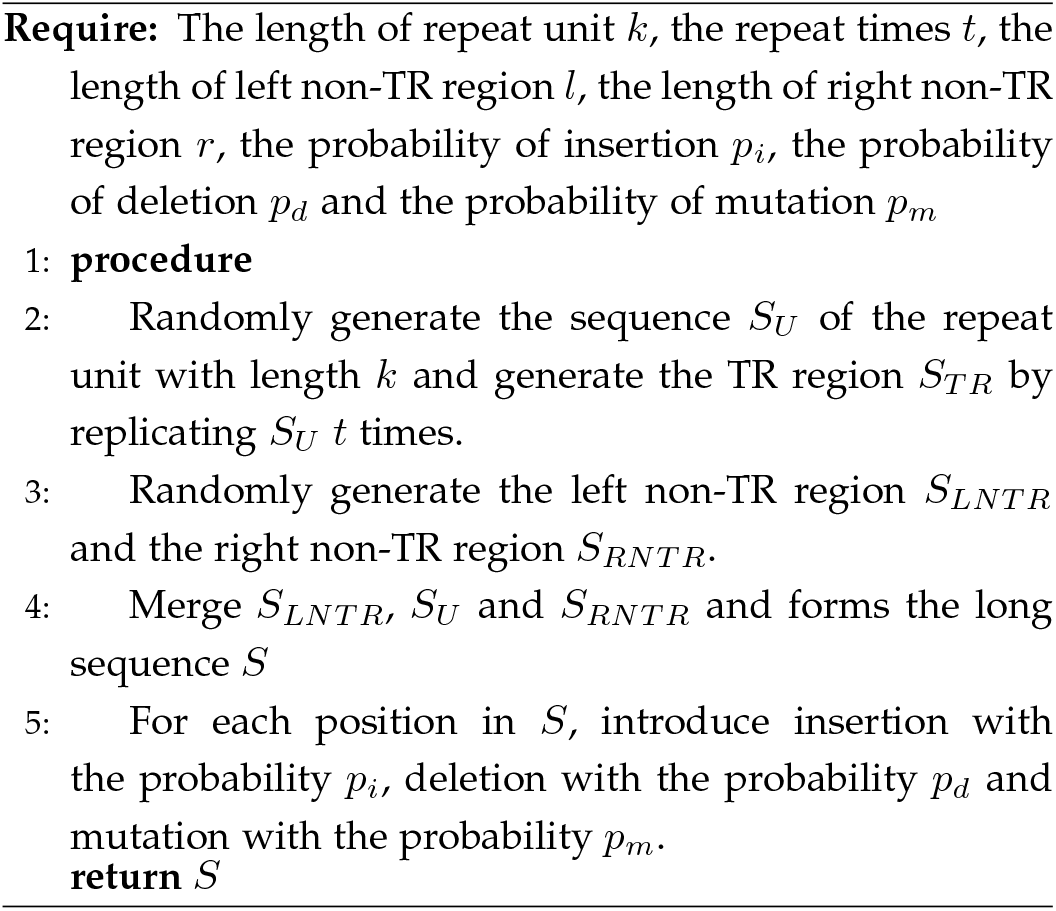

### 3.2 Transformed similarity matrix and deep learning model

After converting each sub-sequence from the previous step into a positional weighted matrix (PWM) with shape (4, *N*) (where *N* is the length of the sequence), we further transformed it into a transformed similarity matrix (TSM) using Algorithm 2. The resulting TSM was used as input for the deep learning model.

DeepTRs comprises 17 individual deep learning models, with each model responsible for predicting a specific k-mer. These models use the TSM corresponding to the k-mer as input. Each model has two convolutional layers for feature extraction and three fully connected layers for classification, each followed by a corresponding ReLU layer. The output of the final layer is the predicted probability that the TSM represents a tandem repeat.

#### Algorithm 2 Modality conversion from PWM to TSM

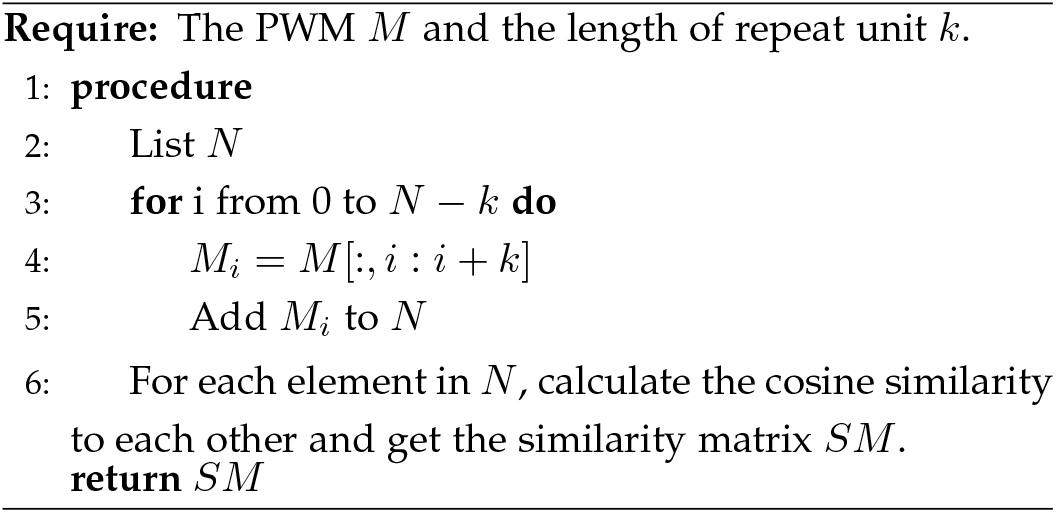

### 3.3 Model training and inference

During model training, the learning rate was set to 1e–5 and the Adam optimizer was used. A workstation with 252 GB RAM, 112 CPU cores and 2 Nvidia V100 GPUs was adopted for all experiments. DeepTRs was developed based on Python3.7, PyTorch1.9.1 and CUDA11.4. A detailed list of dependencies could be found in our code availability.

### 3.4 Evaluation matrices

To assess the performance of models, we adopted metrics including accuracy and macro-F1, which are defined below:

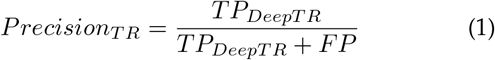

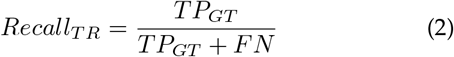

where *TP*_*DeepT R*_ represents the total number of predictions made by DeepTR that are found in the ground truth (GT) dataset, including those that were successfully retrieved from the complete annotations of RepeatMasker. *FP* corresponds to the remaining predictions made by DeepTR. *TP*_*GT*_ denotes the number of GT that match the predictions made by DeepTR, and *FN* represents the number of GT that are not covered by DeepTR’s predictions.

### 3.5 DeepTR Visualizer

DeepTR Visualizer was developed with Plotly 5.14.1 and Dash 2.9.3. As shown in Figure 1, DeepTR Visualizer includes a parameter setting panel and a result visualization panel. The result visualization panel includes the visualisation of PWM, integrated prediction and Voting results.

## 4 Discussion

Traditional TR identification methods are often limited to processing genomes obtained through sequence assembly and cannot directly start detection from sequencing reads. For example, two widely used tools, TRF and RepeatMasker, do not have the capability to directly identify TRs from sequencing reads. Furthermore, the inflexibility of detection mode and parameters hinders the accuracy and completeness of the identification, rendering the results unsatisfactory. For example, TRF’s recognition results can be significantly influenced by its parameters, leading to incomplete detection outcomes. These shortcomings result in existing TR variation identification methods being associated with high computational cost, limited detection sensitivity, precision and comprehensiveness.

The proposed DeepTRs enables direct TR variation identification from raw Nanopore sequencing reads and achieves high sensitivity, accuracy, and completeness results through the multi-modal conversion of Nanopore reads alignment and deep learning. First of all, convert the multiple sequence alignment into an image. The TR regions will appear in an apparently regular color and arrangement on this image. Afterwards, we employ deep learning to acquire and represent these features. Finally, we use these learned features to identify new TRs. The advantages of DeepTRs should be represented in the following aspects: 1) it can perform TR identification directly based on nanopore sequencing raw reads, even under extremely low-depth sequencing conditions (depth*>*=5), which can greatly reduce the detection cost and time complexity. 2) it has high sensitivity and fault tolerance, and overcomes the problem of poor detection integrity of traditional tools caused by inflexible parameters and detection modes. 3) Due to its superior performance in TR detection, DeepTRs offers greater benefits for high-precision and comprehensive TR variation detection. 4) In the future, DeepTRs will offer a more powerful web version that includes an enhanced visual display module and an association analysis module specifically designed for complex diseases.

## 5 Acknowledgements

## Data availability

All described datasets are publicly available through the corresponding repositories. The nanopore reads for sample HG001 can be accessed at https://s3-us-west-2.amazonaws.com/human-pangenomics/NHGRIUCSCpanel/HG001/nanopore/, while the corresponding assembly (GCA 002077035.3) is available at https://www.ncbi.nlm.nih.gov/assembly/GCA002077035.3. The nanopore reads for sample HG002 can be accessed at ftp://ftp-trace.ncbi.nlm.nih.gov/giab/ftp/data/AshkenazimTrio/HG002_NA24385son/UCSCUltralong_OxfordNanopore_Promethion/, while the corresponding assembly (GCA 018852605.1) is available at https://www.ncbi.nlm.nih.gov/data-hub/genome/GCA_018852605.1/. The nanopore reads for sample HG003 can be accessed at ftp://ftp-trace.ncbi.nlm.nih.gov/giab/ftp/data/AshkenazimTrio/HG003NA24149_father/UCSC_Ultralong_OxfordNanopore_Promethion/, while the corresponding assembly (GCA 001549605.1) is available at https://www.ncbi.nlm.nih.gov/assembly/GCA_001549605.1/. The nanopore reads for sample HG004 can be accessed at ftp://ftp-trace.ncbi.nlm.nih.gov/giab/ftp/data/Ashken_azimTrio/HG004_NA24143_mother/UCSC_Ultralong_OxfordNanopore_Promethion/, while the corresponding assembly (GCA 001549595.1) is available at https://www.ncbi.nlm.nih.gov/assembly/GCA001549595.1/. The nanopore reads for sample HG005 can be accessed at ftp://ftp-trace.ncbi.nlm.nih.gov/giab/ftp/data/ChineseTrio/HG005_NA24631_son/UCSCUltralong_OxfordNanopore_Promethion/, while the corresponding assembly (GCA 018506945.1) is available at https://www.ncbi.nlm.nih.gov/assembly/GCA_018506945.1. The nanopore reads for sample HG00733 can be accessed at https://s3-us-west-2.amazonaws.com/human-pan_genomics/index.html?prefix=NHGRI_UCSC_panel/H_G00733/nanopore/, while the corresponding assembly (GCA 002208065.1) is available at https://www.ncbi.nlm.nih.gov/assembly/GCA_002208065.1/. The nanopore reads for nine lung cancer samples (DRR171452, DRR171453, DRR171454, DRR171429, DRR171430, DRR171431, DRR171432, DRR171433, and DRR203145) are are available at https://trace.ncbi.nlm.nih.gov/. The version of human reference genome used in this study is GCF 000001405.39 GRCh38.p13.

## Code availability

The DeepTRs software is publicly available at https://github.com/JoshuaChou2018/DeepTR.

## References

[1] G. A. P. Frank R. Wendt and R. Polimanti, “Phenome-wide association study of loci harboring de novo tandem repeat mutations in uk biobank exomes,” Nature communications, vol. 13, p. 7682, 2022.

[2] X. M. M. M. B. Alexander S. Leonard Danang Crysnanto and H. Pausch, “Graph construction method impacts variation representation and analyses in a bovine super-pangenome,” Genome Biology, vol. 24(1), p. 124, 2023.

[3] R. Y. M. G. Nima Mousavi, Sharona Shleizer-Burko, “Profiling the genome-wide landscape of tandem repeat expansions,” Nucleic acids research, vol. 47(15), pp. e90–e90, 2019.

[4] R. K. Y. Terence Gall-Duncan Nozomu Sato and C. E. Pearson, “Advancing genomic technologies and clinical awareness accelerates discovery of disease-associated tandem repeat sequences,” Genome research, vol. 32(1), pp. 1–27, 2022.

[5] A.-A. R. e. a. Erwin G S, Gürsoy G, “Recurrent repeat expansions in human cancer genomes,” Nature, vol. 613(7942), pp. 96–102, 2023.

[6] M. N.-e. a. Mitra I, Huang B, “Patterns of de novo tandem repeat mutations and their role in autism,” Nature, vol. 589(7841), pp. 246–250, 2021.

[7] K. E.-e. a. Hamanaka K, Yamauchi D, “Genome-wide identification of tandem repeats associated with splicing variation across 49 tissues in humans,” Genome Research, vol. 33(3), pp. 435–447, 2023.

[8] P. R. e. a. Ciobanu C G, Nucă I, “Narrative review: Update on the molecular diagnosis of fragile x syndrome,” International Journal of Molecular Sciences, vol. 24(11), p. 9206, 2023.

[9] M. S. Masutani B, Kawahara R, “Decomposing mosaic tandem repeats accurately from long reads,” Bioinformatics, vol. 39(4), p. btad185, 2023.

[10] B. G., “Tandem repeats finder: a program to analyze dna sequences,” Nucleic acids research, vol. 27(2), pp. 573–580, 1999.

[11] M. R. L. Haley A L, “Transposable element diversity remains high in gigantic genomes,” Journal of Molecular Evolution, vol. 90(5), pp. 332–341, 2022.

[12] S. O. e. a. Delucchi M, Schaper E, “A new census of protein tandem repeats and their relationship with intrinsic disorder,” Genes, vol. 11(4), p. 407, 2020.

[13] K. A. V. Jorda J, “T-reks: identification of tandem repeats in sequences with a k-means based algorithm,” Genes, vol. 25(20), pp. 2632–2638, 2009.

[14] H. L. Y. e. a. Lim K G, Kwoh C K, “Review of tandem repeat search tools: a systematic approach to evaluating algorithmic performance,” Briefings in bioinformatics, vol. 14(1), pp. 67–81, 2013.

[15] A. M. A. M. Mokhtar M M, “Ssrome: an integrated database and pipelines for exploring microsatellites in all organisms,” Nucleic Acids Research, vol. 47(D1), pp. D244–D252, 2019.

[16] S. T. e. a. Osmanli Z, Falgarone T, “The difference in structural states between canonical proteins and their isoforms established by proteome-wide bioinformatics analysis,” Biomolecules, vol. 12(11), p. 1610, 2022.

[17] V. A. Pellegrini M, Renda M E, “Trstalker: an efficient heuristic for finding fuzzy tandem repeats,” Bioinformatics, vol. 26(12), pp. i358–i366, 2010.

[18] V. A. e. a. Avvaru A K, Sharma D, “Msdb: a comprehensive, annotated database of microsatellites,” Nucleic acids research, vol. 48(D1), pp. D155–D159, 2020.

[19] H. A. Bharti P K, “Empirical evaluation of in silico microsatellites mining tools designed using nextgen technology in crops,” 2022 7th International Conference on Computing, Communication and Security (ICCCS), vol. 48(D1), pp. 1–5, 2022.

[20] W.-R. J. e. a. Schoelmerich M C, Sachdeva R, “Tandem repeats in giant archaeal borg elements undergo rapid evolution and create new intrinsically disordered regions in proteins,” Plos Biology, vol. 21(1), p. e3001980, 2023.

[21] T. H. G. S. V. C. Tsung-Yu Lu and M. J. P. Chaisson, “Profiling variable-number tandem repeat variation across populations using repeat-pangenome graphs,” Nature communications, vol. 12(1), p. 4250, 2021.

[22] W. S. e. a. Takeuchi T, Suzuki Y, “A high-quality, haplotype-phased genome reconstruction reveals unexpected haplotype diversity in a pearl oyster,” DNA Research, vol. 29(6), p. dsac035, 2022.

[23] W. L. e. a. Ji Y, Feng S, “Orthologous microsatellites, transposable elements, and dna deletions correlate with generation time and body mass in neoavian birds,” Science Advances, vol. 8(35), p. eabo0099, 2022.

[24] M. E. W. Morishita S, Ichikawa K, “Finding long tandem repeats in long noisy reads,” Bioinformatics, vol. 37(5), pp. 612–621, 2021.

[25] e. a. Pereira, Luísa, “Popaffiliator: online calculator for individual affiliation to a major population group based on 17 autosomal short tandem repeat genotype profile,” International Journal of Legal Medicine, vol. 125, pp. 629–636, 2011.

[26] N. S. F. D. R. N. V. A. V. K. Vladimir Perovic Jeremy Y Leclercq, “Tally-2.0: upgraded validator of tandem repeat detection in protein sequences,” Bioinformatics, vol. 36(10), p. 3260–3262, 2020.

[27] X. I. e. a. Fazal S, Danzi M, “Rexprt: a machine learning tool to predict pathogenicity of tandem repeat loci,” bioRxiv, vol. 2023.03, p. 22.533484, 2023.

[28] B. B. e. a. Sitarčík J, Vinař T, “Warpstr: Determining tandem repeat lengths using raw nanopore signals,” Bioinformatics, vol. 39(6), p. btad388, 2023.

[29] A. M. M. P. G.-A. B. L. D. Li Fang Qian Liu and K. Wang, “Deeprepeat: direct quantification of short tandem repeats on signal data from nanopore sequencing,” Genome biology, vol. 23(1), p. 108, 2022.

[30] P. Guillaume and S. Grudinin, “Deepsymmetry: using 3d convolutional networks for identification of tandem repeats and internal symmetries in protein structures,” Bioinformatics, vol. 35(24), pp. 5113–5120, 2019.

[31] W. Y. S. J. Y. Z. Lang J, Xu Z, “Nanostr: A method for detection of target short tandem repeats based on nanopore sequencing data,” Frontiers in Molecular Biosciences, vol. 10, p. 1093519, 2023.

[32] G. J. e. a. Eraslan G, Avsec Z?, “Deep learning: new computational modelling techniques for genomics,” Nature Reviews Genetics, vol. 20(7), pp. 389–403, 2019.

[33] L. V. V. e. a. Mishra V, Re D B, “Systematic elucidation of neuron-astrocyte interaction in models of amyotrophic lateral sclerosis using multi-modal integrated bioinformatics workflow,” Nature Reviews Genetics, vol. 11(1), p. 5579, 2020.

[34] L. J. e. a. Huang K, Lin B, “Predicting colorectal cancer tumor mutational burden from histopathological images and clinical information using multi-modal deep learning,” Bioinformatics, vol. 38(22), pp. 5108–5115, 2022.

[35] R. E. e. a. Giesselmann P, Brändl B, “Analysis of short tandem repeat expansions and their methylation state with nanopore sequencing,” Nature Biotechnology, vol. 37(12), pp. 1478–1481, 2019.

[36] I. K. e. a. Dolzhenko E, Weisburd B, “Reviewer: haplotype-resolved visualization of read alignments in and around tandem repeats,” Genome medicine, vol. 14(1), p. 84, 2022.

[37] e. a. Dolzhenko Egor, “Expansionhunter: a sequence-graph-based tool to analyze variation in short tandem repeat regions,” Bioinformatics, vol. 35(22), pp. 4754–4756, 2019.

[38] e. a. Dolzhenko Egor, “Expansionhunter denovo: a computational method for locating known and novel repeat expansions in short-read sequencing data,” Genome biology, vol. 21, pp. 1–4, 2020.

[39] e. a. Dashnow, Harriet, “Stretch: detecting and discovering pathogenic short tandem repeat expansions,” Genome biology, vol. 19(1), pp. 1–13, 2018.

[40] M. T. e. a. Mitsuhashi S, Frith M C, “Tandem-genotypes: robust detection of tandem repeat expansions from long dna reads,” Genome biology, vol. 20, pp. 1–17, 2019.

[41] B. L. e. a. De Roeck A, De Coster W, “Nanosatellite: accurate characterization of expanded tandem repeat length and sequence through whole genome long-read sequencing on promethion,” Genome biology, vol. 20, pp. 1–16, 2019.

[42] B. V. e. a. Bolognini D, Magi A, “Tricolor: tandem repeat profiling using whole-genome long-read sequencing data,” Gigascience, vol. 9(10), p. giaa101, 2020.

[43] F. J. M. e. a. Chiu R, Rajan-Babu I S, “Straglr: discovering and genotyping tandem repeat expansions using whole genome longread sequences,” Genome biology, vol. 21(1), p. 224, 2021.

[44] L. H, “New strategies to improve minimap2 alignment accuracy,” Bioinformatics, vol. 37, pp. 4572–4574, 2021.

[45] W. A. F. T. R. J. H. N. M. G. A. G. D. R.. G. P. D. P. S. Li H, Handsaker B, “The sequence alignment/map format and samtools,” Bioinformatics, vol. 25(16), pp. 2078–9, 2009.

